# Ectodomain shedding of L-selectin by ADAM17 in canine neutrophils

**DOI:** 10.1101/2020.04.06.027433

**Authors:** Kristin M. Snyder, Camille McAloney, Joshua S. Montel, Jaime F. Modiano, Bruce Walcheck

**Author notes:** Corresponding author: Dr. Bruce Walcheck. These authors contributed equally to the work (Kristin M. Snyder, Camille McAloney).

## Abstract

The adhesion protein L-selectin (CD62L) is expressed at high levels by circulating neutrophils and has a critical role in initiating their recruitment at sites of inflammation. L-selectin expression is rapidly downregulated upon neutrophil activation by various stimuli through a proteolytic process referred to as ectodomain shedding, which regulates L-selectin’s binding avidity and neutrophil recruitment. In humans and mice, L-selectin shedding is primarily mediated by ADAM17 (a disintegrin and metalloproteinase 17). L-selectin expression is also rapidly downregulated by canine neutrophils upon their activation; however, the role of ADAM17 in this process has not been previously investigated. We show that a highly selective inhibitor of ADAM17, but not an inhibitor of its most closely related family member ADAM10, effectively blocked L-selectin downregulation from the surface of canine neutrophils following their activation. The ADAM17 inhibitors did not block the rapid upregulation of the CD18 integrin Mac-1 (CD11b/CD18), showing that they did not broadly impair neutrophil activation. To directly examine the expression of ADAM17, we used several anti-human ADAM17 mAbs. Many did not stain canine neutrophils; however, the ADAM17 function-blocking mAbs MEDI3622 and D1(A12) did stain and also blocked L-selectin downregulation. Taken together, our findings provide the first direct evidence that ADAM17 is a primary sheddase of canine L-selectin, which may serve as an important therapeutic target to prevent neutrophil dysfunction during conditions of excessive inflammation, such as sepsis.

## Introduction

L-selectin (CD62L), where “L” signifies leukocyte, is a member of the Selectin family of adhesion proteins [1, 2]. L-selectin facilitates the accumulation of neutrophils along the vascular endothelium, the initial step of a “multi-step” process of their migration from the blood vessel lumen, through the vascular wall, and into the underlying tissue at sites of inflammation [3]. L-selectin is expressed at high levels by resting neutrophils and is primarily distributed on the ends of cellular microvilli [4]. This distribution pattern facilitates their capture and rolling along the vascular wall under hydrodynamic shear stress [3]. In addition to its role in the direct attachment of free-flowing neutrophils to the vascular endothelium, L-selectin also mediates neutrophil attachment to previously tethered neutrophils, and this process of indirect capture can greatly amplify neutrophil accumulation [5-7].

Unlike most adhesion molecules expressed by neutrophils, L-selectin undergoes a process referred to as ectodomain shedding. This proteolytic event occurs upon cell activation and results in a rapid downregulation of L-selectin expression from the cell surface [1, 2, 8]. L-selectin shedding, in part, regulates the avidity of its interactions with vascular ligands and neutrophil tethering efficiency [9]. However, excessive L-selectin shedding by circulating neutrophils has been shown to occur during excessive sterile inflammation and infection, which was associated with their diminished recruitment at sites of infection and increased bacterial levels and spread [10, 11].

ADAM17, a member of the <u>a d</u>isintegrin <u>a</u>nd <u>m</u>etalloproteinase family, is a type-1 transmembrane protein that typically cleaves its substrates in a *cis* manner [1]. Human and mouse L-selectin are well described ADAM17 substrates, which are cleaved at a single extracellular region proximal to the cell membrane [12-15]. L-selectin does not undergo shedding from ADAM17-null neutrophils when activated *in vitro*, and this process is markedly impaired *in vivo* in ADAM17 gene-targeted mice during sterile inflammation and bacterial infection [10, 13, 16]. However, over-expression of ADAM10, the most similar family member to ADAM17 in terms of amino acid sequence and structure [17], results in low level L-selectin shedding in transfected cells, indicating its potential redundant role in L-selectin shedding [18].

Canine L-selectin has been characterized at the protein, cellular, and functional levels and is very similar to human L-selectin [19, 20]. Canine L-selectin undergoes rapid downregulation in expression upon neutrophil activation, suggesting it is similarly regulated by ectodomain shedding [19]. At this time, the role of ADAM17 in L-selectin shedding by canine neutrophils has not been determined. We demonstrate ADAM17 expression by canine neutrophils and its role in downregulating L-selectin expression upon neutrophil stimulation with various stimuli. Given that ADAM17 is a central regulator of inflammation and neutrophil effector functions in humans [1], ADAM17 may provide a critical therapeutic target to prevent neutrophil dysfunction in dogs during inflammatory disorders, such as sepsis.

## Materials and methods

### Animals

Peripheral blood was collected from healthy dogs. Blood collection was carried out in strict accordance with the recommendations in the Guide for the Care and Use of Laboratory Animals of the National Institutes of Health. The protocol was approved by the Institutional Animal Care and Use Committee of the University of Minnesota (Protocol Number: 21903-36913A).

### Antibodies

D1(A12) (MilliporeSigma, St. Louis, MO) is an anti-human ADAM17 mAbs and has been shown to block its proteolytic activity [21]. MEDI3622 is an anti-human ADAM17 mAb and has been shown to that block its proteolytic activity [22, 23]. For this study, MEDI3622 was generated as a human IgG1 by Syd Labs (Natick, MA) using its variable heavy and variable light chain sequences. The anti-human ADAM17 mAbs 111633 and 111623 were purchased from R&D Systems (Minneapolis, MN). The mouse anti-human ADAM17 mAb M220 (generously provided by Dr. R. Black, Immunex, Seattle, WA) has been previously described [24]. The mAb CL2/6 (Thermo Fisher Scientific, Waltham, MA) recognizes human E-selectin and canine L-selectin [19]. The mAb R15.7 (generously provided by Dr. R. Rothlein, Boehringer Ingelheim, Ridgefield, CT) has been previously shown to recognize canine CD18 [25]. Isotype-matched negative control mouse and human antibodies were purchased from R&D Systems and MilliporeSigma, respectively. Allophycocyanin-conjugated F(ab′)_2_ goat anti-mouse or donkey anti-human IgG (H+L) secondary antibodies were purchased from Jackson ImmunoResearch Laboratories (West Grove, PA).

### Canine neutrophil isolation and treatment

Canine peripheral blood was collected from healthy donors in K_2_-EDTA blood collection tubes (Becton, Dickinson and Company, Franklin Lakes, NJ). Whole blood was diluted ten-fold with red blood cell lysis buffer (ammonium chloride, 155 mM; potassium bicarbonate, 10 mM; and disodium EDTA, 0.1 mM) and incubated for 10 minutes at room temperature. Leukocytes were washed thoroughly in PBS buffer (without Ca^+2^ and Mg^+2^) (Lonza, Walkersville, MD) and cell viabilities were assessed using trypan blue exclusion. Leukocytes (2.5×10^6^/ml in PBS) were stimulated with 1-15ng/mL phorbol-12-myristate-13-acetate (PMA) (MilliporeSigma), 1µg/mL formyl peptide receptor-like 1 agonist (MilliporeSigma), or 10µg/mL *Pseudomonas* lipopolysaccharide (LPS) (MilliporeSigma), as we have previously described [26]. Cells were stimulated for 30 minutes at 37°C, which was stopped by extensive cell washing with PBS at 4°C. Some cells were pre-incubated for 30 minutes on ice with ADAM10 and ADAM17 hydroxymate-based small molecule inhibitors or function blocking antibodies prior to activation. The selective ADAM17 inhibitor BMS566394 (MedKoo Biosciences, Morrisville, NC), referred to as inhibitor 32 in reference [27], was used at 5 μM, as we have previously described [28]; the selective ADAM10 inhibitor GI254023X (R&D Systems) is 10-fold more selective for ADAM10 than ADAM17 in cellular assays and preferentially blocks ADAM10 in cells at a concentration ranging from 0.2-1 μM [29, 30]. GI254023X was used at 0.5 μM, as we have previously described [28]. INCB3619 (MedKoo Biosciences) blocks the activity of ADAM10 and ADAM17 [31]. INCB3619 was used at 10µM, as we have previously described [28]. All inhibitors were resuspended in DMSO and used at the indicated concentrations. The anti-ADAM17 humanized monoclonal antibodies MEDI3622 and D1(A12) were used at 10 μg/ml.

### Cell staining and flow cytometric analysis

For cell staining, Fc receptor and nonspecific antibody binding sites were blocked for 30 minutes using 25% canine serum and 25% FBS in PBS prior to their staining with antibodies. All cell staining was analyzed on FACSCanto or FACSCelesta instruments (BD Biosciences, San Jose, CA). Canine neutrophils were identified based on their forward and side light-scattering characteristics [32]. Comparison between two groups was done by a Student’s t test.

### Pairwise Sequence Alignment

To identify the MEDI3622 binding site for ADAM17 across several species, amino acid sequences of human, mouse, rat, canine (predicted), feline (predicted), and rabbit (predicted) ADAM17 were retrieved from UniProtKB. Amino acid sequences were aligned using Clustal Omega (Dublin, Ireland).

## Results

### Effects of selective ADAM10 and ADAM17 inhibitors on L-selectin downregulation following canine neutrophil activation

Others have reported that canine neutrophils rapidly downregulate L-selectin upon their activation with various stimuli [19]. For instance, overt activation of canine neutrophils with PMA resulted in the rapid downregulation of L-selectin expression, as determined by flow cytometry (**Fig 1**). To establish whether ADAM10 and/or ADAM17 were involved in this process, we utilized several small molecule hydroxamate-based metalloproteinase inhibitors. INCB3619 inhibits both ADAM10 and ADAM17 [31]. We found that INCB3619 effectively blocked L-selectin downregulation following neutrophil activation with PMA (**Fig 1A**). To evaluate the role of ADAM10 in this process, we used the selective inhibitor GI254023X [29]. This inhibitor, however, did not significantly affect L-selectin downregulation following neutrophil activation (**Fig 1A**). The hydroxamate BMS566394 has a potency that is orders of magnitude higher for ADAM17 than ADAM10 [27], and we found that it effectively blocked L-selectin downregulation upon neutrophil activation at a level equivalent to INCB3619 (**Fig 1A**). Downregulation of L-selection was also observed upon neutrophil activation via the appropriate pattern recognition receptors using a formyl peptide or LPS (**Fig 1B**). BMS566394 also blocked L-selectin downregulation by canine neutrophils activated by these stimuli (**Fig 1B**). Our findings suggest a primary role for ADAM17 in shedding canine L-selectin. Of interest is that the membrane proximal region at which ADAM17 cleaves human L-selectin is 100% homologous and 87.5% identical at the amino acid level to the membrane proximal region of canine L-selectin, indicating that canine L-selectin may be cut at the same cleavage site (**Fig 1C**).

**Fig 1.**
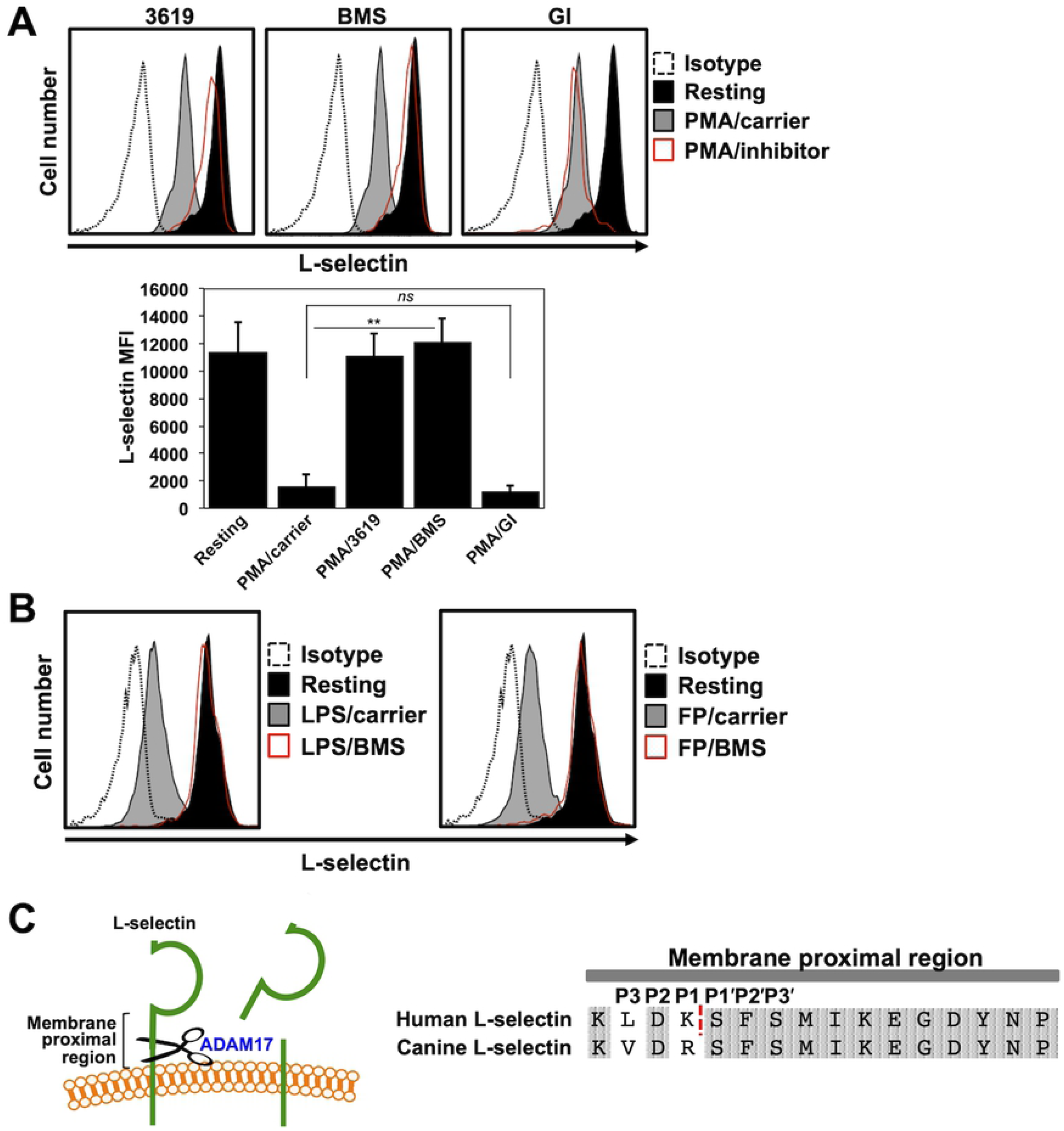
ADAM17 activity in canine neutrophils. **A**. Canine peripheral blood leukocytes were treated with or without PMA in the presence or absence of the ADAM10 and ADAM17 inhibitor INCB3619 (3619), the ADAM17 inhibitor BMS566394 (BMS), the ADAM10 inhibitor GI254023X (GI), or DMSO carrier for 30 minutes at 37°C, as indicated. The leukocytes were then stained with an anti-L-selectin mAb or an appropriate isotype negative control antibody and assessed by flow cytometry. Neutrophils were gated based on their characteristic forward and side light scatters. All histograms show representative data and the bar graphs show mean ± SD with at least 3 animals per treatment. The x-axis of all histograms = Log 10 fluorescence. Statistical significance is indicated as **p < 0.01 PMA/carrier vs. PMA/3619 and PMA/BMS. PMA/carrier vs. PMA/GI was not significant (*ns*). MFI = Mean fluorescence intensity. **B**. Leukocytes were treated with the ADAM17 inhibitor BMS as described above; however, LPS or formyl peptide (FP) were used as stimuli. **C.** L-selectin undergoes ectodomain shedding by ADAM17 within a membrane proximal region, as indicated. Alignment of the extracellular membrane proximal regions of human and canine L-selectin. The dashed line indicates the site of cleavage in human L-selectin, as previously described [14]. The amino acid sequences of human and canine L-selectin are from the NCBI reference sequences NM_000655.4 and XM_537201.4, respectively.

The CD18 integrins, including Mac-1 (CD11b, CD18), are additional adhesion molecules expressed on neutrophils that are important for their attachment and transmigration through the vascular endothelium [3]. In contrast to L-selectin, Mac-1 surface expression is rapidly upregulated following neutrophil activation, as the adhesion molecule resides in intracellular granules that are translocated to the cell surface [33]. We found that neither INCB3619, GI254023X, nor BMS566394 treatment of canine neutrophils affect an increase in CD18 expression upon their activation (**Fig 2**), demonstrating that INCB3619 and BMS566394 did impair neutrophil activation as a means of blocking L-selectin down-regulation.

**Fig 2.**
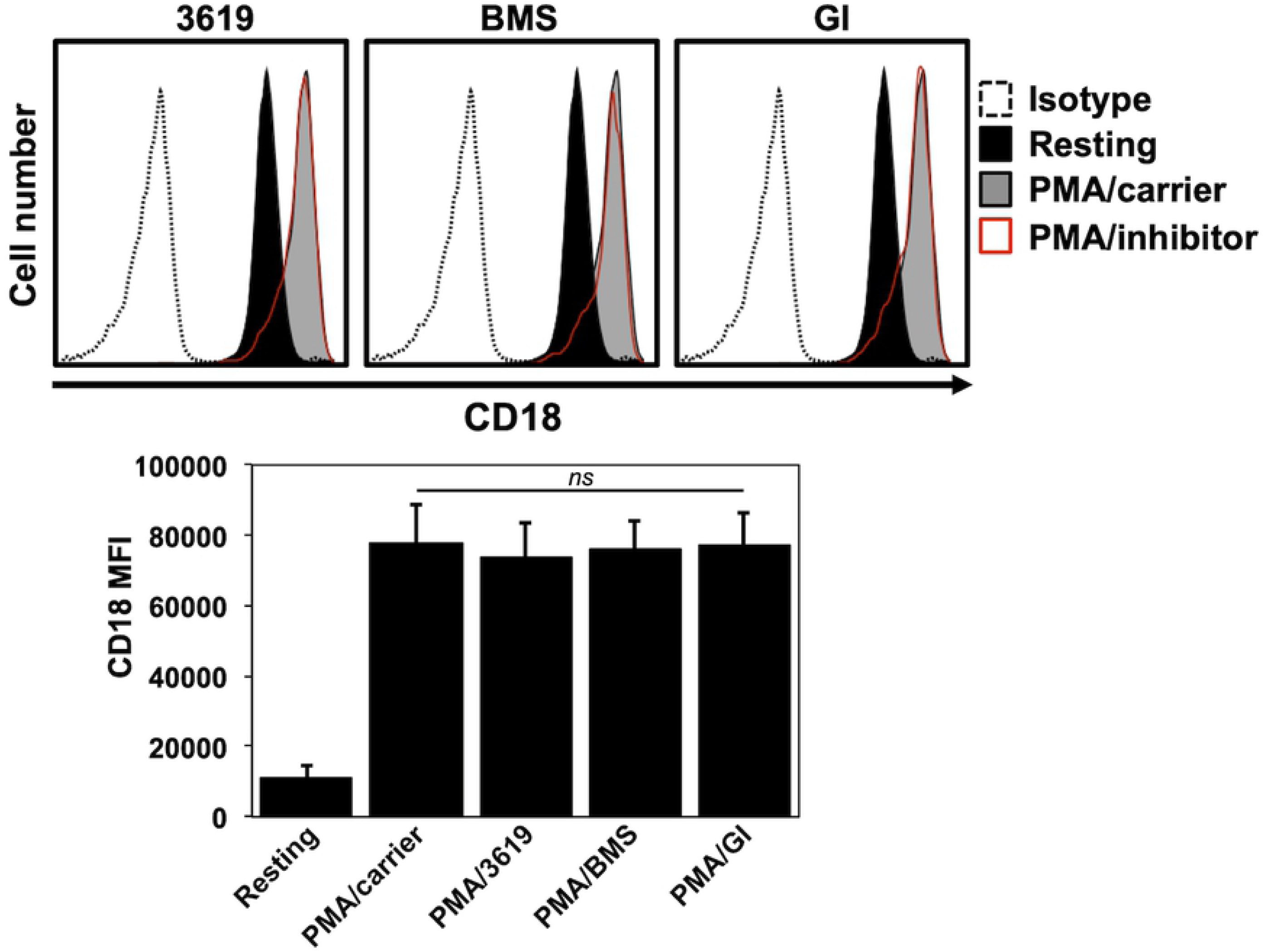
ADAM10 and ADAM17 inhibitors do not affect Mac-1 up-regulation. Peripheral blood leukocytes were treated with or without PMA in the presence or absence of ADAM10 and/or ADAM17 inhibitors, as described in Figure 1. Relative cell-staining levels of CD18 were determined by flow cytometry. Neutrophils were gated based on their characteristic forward and side light scatters. The x-axis of all histograms = Log 10 fluorescence. All histograms show representative data and the bar graphs show mean ± SD with at least 3 animals per treatment.

### ADAM17 expression and function by canine neutrophils

To date, antibodies that detect canine ADAM17 have not been described. The predicted amino acid sequence of canine ADAM17 based on cDNA sequence reveals a 96% identity to human ADAM17 [34]. Therefore, we examined the reactivity of several anti-human ADAM17 mAbs with canine neutrophils. We have previously demonstrated that the anti-ADAM17 mAbs 111633, 111623, and m220 all stained ADAM17 on human cells by flow cytometry [30, 35-37], and in **Figure 3A**, we show that these mAbs stained human peripheral blood neutrophils above that of an isotype-matched negative control antibody, whereas this was not the case for canine peripheral blood neutrophils (**Fig 3B**).

**Fig 3.**
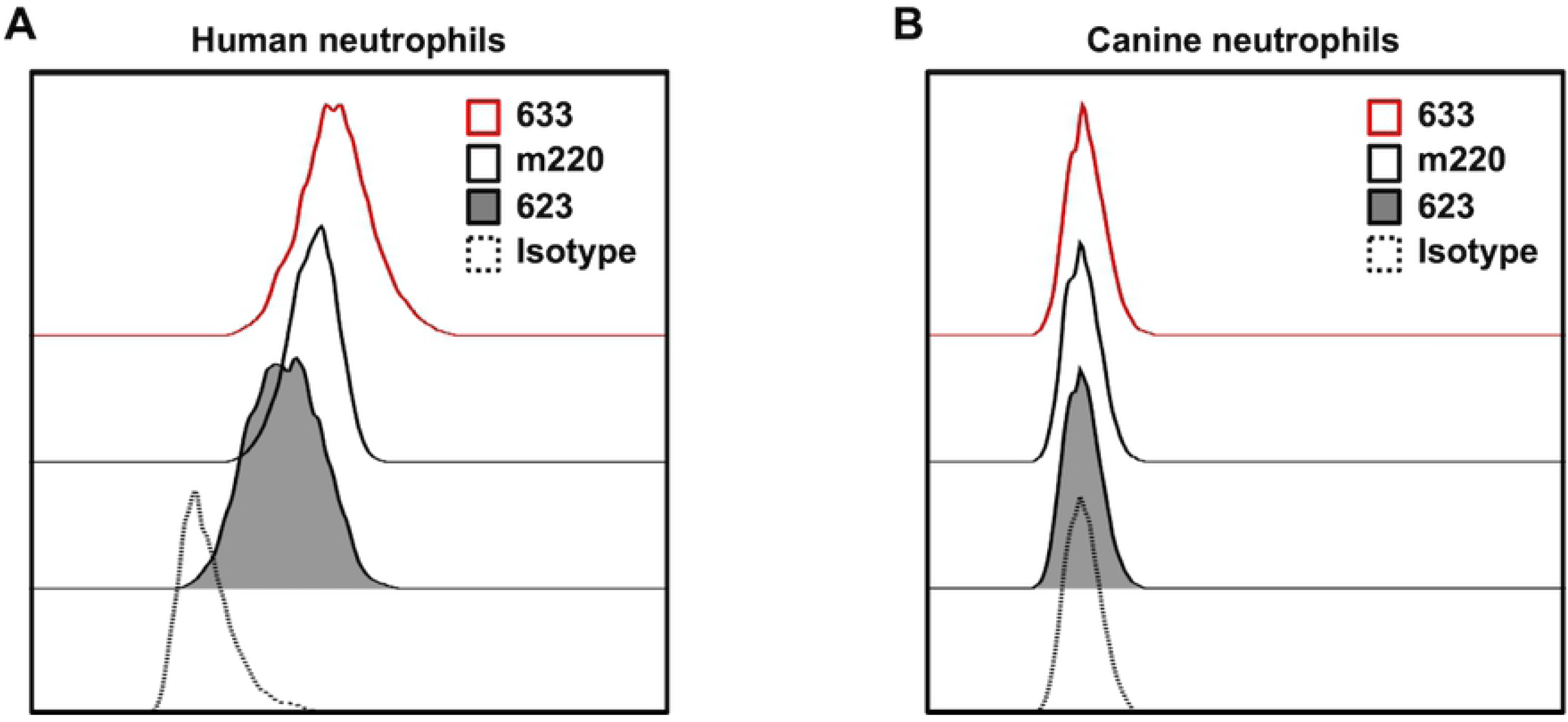
Anti-ADAM17 mAbs stain human but not canine neutrophils. Human (**A**) and canine (**B**) peripheral blood leukocytes were stained by different anti-ADAM17 mAbs and examined by flow cytometry, as indicated. Neutrophils were gated based on their characteristic forward and side light scatters. The x-axis of all histograms = Log 10 fluorescence. Data is representative of at least 3 independent experiments using leukocytes from separate human and canine donors.

MEDI3622 is a well described anti-ADAM17 function blocking mAb [22], which has been shown to block the downregulation of L-selectin on human leukocytes [38]. Its epitope has been mapped to a surface loop located within the catalytic domain of ADAM17 but no other ADAM family members [23]. This region of ADAM17 is highly conserved and MEDI3622 has been previously shown to also recognize mouse ADAM17 [22, 38]. The MEDI3622 epitope occurs between the amino acids P^366^-N^381^ of human ADAM17 [23]. Alignment of this region with the predicted amino acid sequence for canine ADAM17 shows it to be nearly identical (**Fig 4A**). Furthermore, this region of ADAM17 is conserved in several other species, including mouse, rat, feline and rabbit (**Fig 4A**). Consistent with this, MEDI3622 was found to specifically stain canine neutrophils (**Fig 4B**). MEDI3622 also blocked the downregulation of surface L-selectin expression upon canine neutrophil activation (**Fig 4C**). D1(A12) is another anti-human ADAM17 mAb that blocks function [21]. We found this mAb also stained canine neutrophils and blocked the downregulation of L-selectin upon their activation (**Fig 4D, E)**, though not as effectively as MEDI3622 (**Fig 4C**).

**Fig 4.**
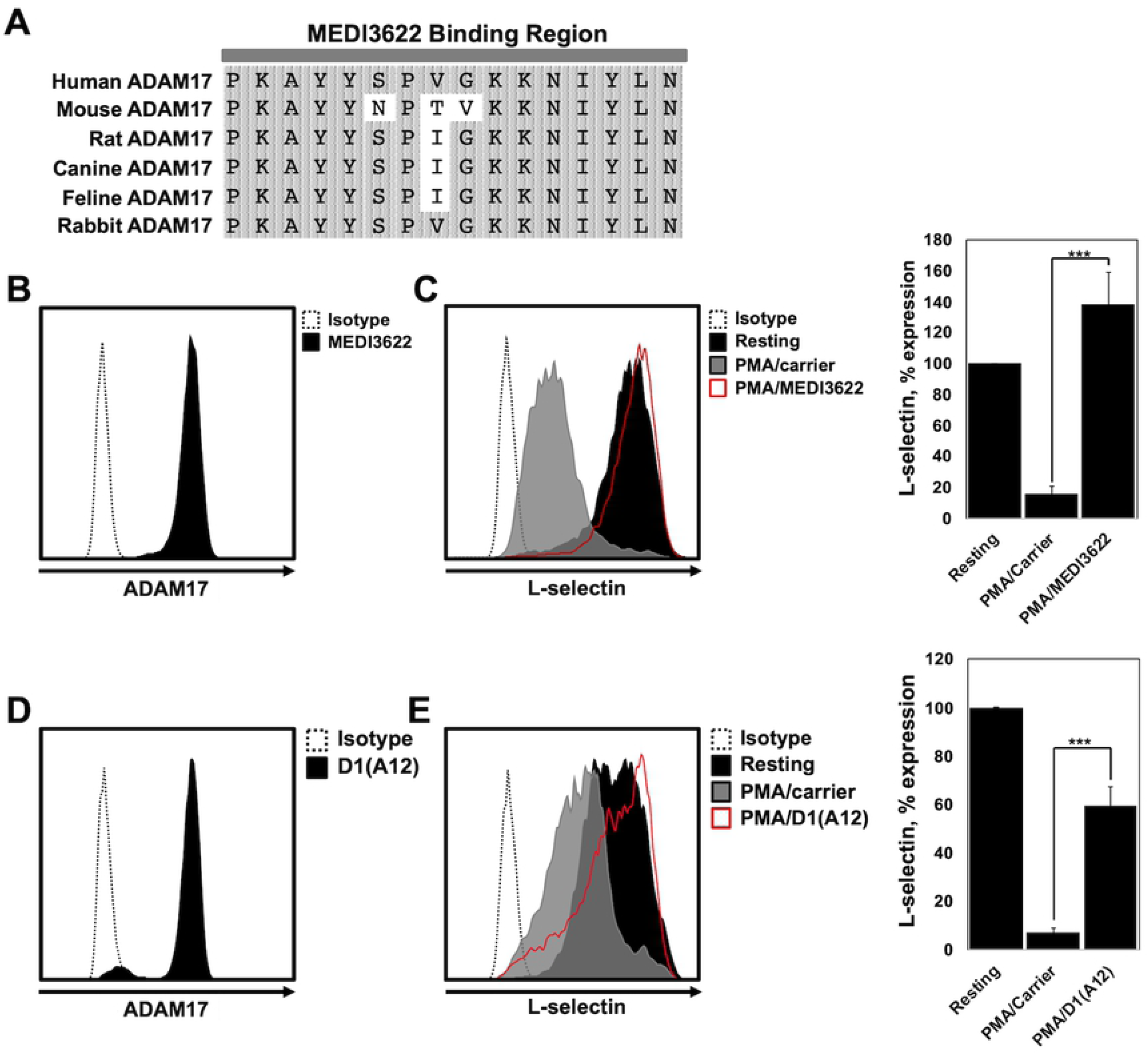
ADAM17 mAbs block the function of canine ADAM17. **A**. Alignment of amino acid sequences for ADAM17 across several species shows a high level of conservation in the MEDI3622 binding site, which occurs between amino acids P^366^-N^381^. Primary accession number (UniProt) for amino acid sequences are as follows: human (P78536), mouse (Q9Z0F8), rat (Q9Z1K9), canine predicted (M1V4E8), feline predicted (M3W8I1), and rabbit predicted (G1SZC8). **B**. Peripheral blood neutrophils were stained with MEDI3622 or an IgG1 isotype negative control antibody and assessed by flow cytometry. **C**. Peripheral blood neutrophils were treated with or without PMA in the presence of MEDI3622. L-selectin shedding was determined via flow cytometry. **D**. Peripheral blood neutrophils were stained with D1(A12) or an IgG1 isotype negative control antibody and assessed by flow cytometry. **E.** Peripheral blood neutrophils were treated with or without PMA in the presence of D1(A12). L-selectin shedding was determined via flow cytometry. The x-axis of all histograms = Log 10 fluorescence. Data is representative of at least 3 independent experiments using leukocytes from separate animals. The bar graph shows mean ± SD with at least 3 animals per treatment. Statistical significance is indicated as ***p <0.001 PMA/carrier vs. PMA/MEDI3622 or PMA/carrier vs PMA/D1(A12).

## Discussion

Here, we demonstrate the function of ADAM17 in canine neutrophils and its role in the downregulation of L-selectin upon their activation. We show that selective small molecule inhibitors of ADAM17, but not ADAM10, as well as the function blocking ADAM17 mAbs MEDI3622 and D1(A12) block canine L-selectin downregulation. The ADAM17 inhibitors did not have any effect on the upregulation of the adhesion protein Mac-1 upon neutrophil activation, indicating that these treatments did not broadly impair neutrophil activation.

Canine and human ADAM17 are highly homologous with 96% sequence identity at the amino acid level [34]. Interestingly is that several anti-human ADAM17 antibodies did not stain canine neutrophils by flow cytometry. In contrast, the anti-ADAM17 mAb MEDI3622 did stain canine neutrophils and also inhibited the sheddase’s activity. Its epitope has been mapped to a loop structure in the protease’s catalytic region that consists of amino acids P^366^-N^381^, which is absent in other ADAM family members [23]. Alignment of this site with the putative sequence for canine ADAM17 revealed high conservation. MEDI3622 also recognizes mouse ADAM17 [22, 38], which has less similarity to human ADAM17 than canine ADAM17 (**Fig 4A**). An alignment of this region from the putative amino acid sequence of ADAM17 from various divergent animal species reveals a high level of conservation (**Fig 4A**), suggesting the likelihood of broad cross-reactivity by MEDI3622. The anti-human ADAM17 mAb D1(A12) also stained canine neutrophils. Interestingly, the variable heavy and light chains of this engineered mAb were designed to recognize distinct epitopes in different regions of ADAM17 [21]. To our knowledge, the specific locations of these epitopes have not been mapped. D1(A12) has been reported not to recognize mouse ADAM17 [21], indicating less cross-reactivity than MEDI3622. D1(A12) also did not block L-selectin downregulation by canine neutrophils as effectively as MEDI3622. Perhaps this is the result of either the heavy or light chain of the mAb not recognizing its respective epitope. However, though both antibodies were used at saturating concentrations, a comprehensive comparison of the inhibitory activity of D1(A12) and MEDI3622 was not performed.

At this time, it is unclear whether canine L-selectin is cleaved by ADAM17 at the same location as human L-selectin. This would appear to be likely considering that the membrane-proximal region of human L-selectin where cleavage occurs is highly conserved in canine L-selectin. L-selectin shedding is thought to be important for controlling neutrophil accumulation levels on the vascular endothelium at sites of inflammation [1, 2]. Considering that L-selectin directs neutrophils to broad sites of inflammation and facilitates their rapid accumulation along the vascular wall through direct and indirect processes, its shedding could provide a means of swiftly curtailing L-selectin adhesion events to prevent an overwhelming influx of these potentially destructive cells. However, excessive or prolonged induction of ADAM17 activity and L-selectin shedding has been reported to impair circulating neutrophils from entering sites of infection in animal models of sepsis and in septic patients [10, 11, 39, 40]. Neutrophil dysfunction is also a characteristic of the early stages of sepsis in dogs [41]. In consideration of this and our findings, ADAM17 may provide a key therapeutic target to maintain neutrophil recruitment during sepsis for improved bacterial clearance. Additionally, ADAM17 is a regulatory checkpoint of other key modulators of neutrophil effector functions and inflammation, including CD16 (FcγRIII) and TNFα [1], and therefore its targeting may also have much broader benefits during damaging inflammation.

## Acknowledgments

We would like to thank Kathy Stuebner and Amber Winter from the University of Minnesota, College of Veterinary Medicine, Clinical Investigation Center for assistance in acquiring canine peripheral blood samples. We also thank Jianming Wu for his assistance with Figure 1C and copy editing the manuscript.

